# An RNA interference approach for functional studies in the sea urchin and its use in analysis of Nodal signaling gradients

**DOI:** 10.1101/2024.06.20.599930

**Authors:** Keen Wilson, Carl Manner, Esther Miranda, Alejandro Berrio, Gregory A Wray, David R McClay

## Abstract

Dicer substrate interfering RNAs (DsiRNAs) destroy targeted transcripts using the RNA-Induced Silencing Complex (RISC) through a process called RNA interference (RNAi). This process is ubiquitous among eukaryotes. Here we report the utility of DsiRNA in embryos of the sea urchin *Lytechinus variagatus (Lv).* Specific knockdowns phenocopy known morpholino and inhibitor knockdowns, and DsiRNA offers a useful alternative to morpholinos. Methods for designing and obtaining specific DsiRNAs that lead to destruction of targeted mRNA are described. DsiRNAs directed against *pks1*, an enzyme necessary for pigment production, show how successful DsiRNA perturbations are monitored by RNA *in situ* analysis and by qPCR to determine relative destruction of targeted mRNA. DsiRNA-based knockdowns phenocopy morpholino- and drug-based inhibition of *nodal* and *lefty*. Other knockdowns demonstrate that the RISC operates early in development as well as on genes that are first transcribed hours after gastrulation is completed. Thus, DsiRNAs effectively mediate destruction of targeted mRNA in the sea urchin embryo. The approach offers significant advantages over other widely used methods in the urchin in terms of cost, and ease of procurement, and offers sizeable experimental advantages in terms of ease of handling, injection, and knockdown validation.

**Highlights:** DsiRNA provides an RNAi approach for perturbation of sea urchin embryos. A dilution series of DsiRNA oligos reveals properties of the Nodal gradient in establishing the Dorsal-Ventral axis.

## Introduction

Knowledge of the gene regulatory mechanisms governing development has greatly expanded over the last two decades largely due to targeted perturbation of gene expression in model developmental systems. Model systems that utilize forward genetics as a tool, as well as reverse genetic approaches, often using RNAi, played a leading role in these advances. Sea urchins became powerful model for the study of early development by leveraging morpholinos in perturbation approaches and the use of reporter constructs for detailed cis-regulatory analyses.

Those efforts uncovered an extensive gene regulatory network (GRN) governing early specification and morphogenetic events up to the larval stage (Davidson et al., 2002; Davidson and Erwin, 2006; McClay, 2011; Peter and Davidson, 2015; Martik et al., 2016; McClay et al., 2021; Massri et al., 2023). The translation blocking morpholino antisense oligos (MASOs) were the workhorse reagents for those discoveries, and MASOs continue to be a widely used method of gene knockdown in the developing sea urchin embryo (Summerton and Weller, 1997; Heasman et al., 2000; Howard et al., 2001; Heasman, 2002; Yaguchi et al., 2022), but like all perturbation approaches there are challenges. Some MASOs easily precipitate out of solution, clog injection needles and make it a challenge to repeat experiments because effective concentrations of a MASO stock solution change over time. Most MASO experiments are “blind” in that the experimenter doesn’t know the level of the MASO-induced reduction in translation. Ideally, MASOs completely block the translation of an mRNA, but unless antibodies are available or splice-blocking MASOs are used, the effectiveness and specificity of the block is unknown. The current “gold standards” for validation of a MASO-based knockdown are rescue of the translation block by expression of a construct that is unaffected by the MASO, and the use of multiple MASOs to knock down the same gene. Rescue by overexpression generally recapitulates neither the tightly controlled spatiotemporal patterns of expression of the endogenous gene, nor the endogenous level of activity, complicating interpretation of the rescue. The reasoning behind two MASOs is that if both MASOs have the same phenotypic response it is more likely attributable to an on-target effect than spurious off-target effects. Published concerns suggest that off-target effects of MASOs can lead to incorrect interpretations (Schulte-Merker and Stainier, 2014). To be fair, morpholinos are not the only perturbation approach that is subject to possible off-target effects. To a greater or lesser degree, all perturbation approaches, including mutations, have the potential for hidden or misleading effects. To counter these concerns each animal system benefits from orthogonal methods to cross validate an experiment.

Recent studies have enhanced the toolkit available for genetic perturbation in the developing sea urchin embryo. CRISPR/Cas9 technology for genome modification (Lin and Su, 2016) has been increasingly used in sea urchins with positive results, including the first instances of directed mutagenesis of adult sea urchins and successful rearing of F2 mutagenized animals (Oulhen and Wessel, 2016, Liu et al, 2019, Yaguchi et al, 2020). While a powerful tool, CRISPR/Cas9 is not without its challenges. Problems of penetrance, off target effects, high reagent cost, mosaicism, and lack of spatiotemporal control are barriers to wide deployment of this technology for use in the developing sea urchin. Other recent additions of tools for understanding gene function include conditional knockdowns with photo-caged MASOs (Bardhan et al., 2021), and a Tet-On system for conditional control of expression (Khor and Ettensohn, 2023). Each of these advances expands the diversity of experimental possibility, and the capacity to validate results via independent approaches.

One technology that has not been realized in sea urchins is RNA interference (RNAi). Since 1990 when RNAi-mediated knockdown began (Napoli et al., 1990; Fire et al., 1991; Lee et al., 1993; Guo 1995), and double stranded RNA was found to be responsible (Fire et al., 1998), the use of RNAi became, and remains today, a valuable tool for functional studies in many model embryo systems. In the sea urchin, however, the successful adaptation of MASOs for perturbations (Howard et al, 2001) occurred at about the same time RNAi became widely adopted. As a result, the pursuit of RNAi-based gene knockdown technologies was largely put aside in favor of MASO technology.

Since then, much has been learned about the mechanism of action and the molecular components of RNAi (Hammond et al., 2001; Sharp, 2001; Agrawal et al., 2003). RNAi is induced on detection of dsRNA by Dicer, which cleaves the dsRNA into 21-26 bp pieces (Zamore et al., 2000; Hannon, 2002; Carmell and Hannon, 2004) called small interfering RNAs (siRNAs). Dicer-bound RNA then associates with Argonaut, TRBP, and other proteins to form the RNA-Induced Silencing Complex (RISC) (Chendrimada et al., 2005). A single strand from the siRNA guides the RISC complex by sequence complementarity to target mRNAs, where the RISC mediates the mRNA fragmentation. The machinery responsible for RNAi is ubiquitous and ancient, present in archaea, fungi, plants, and animals (Shabalina and Koonin, 2008), including sea urchins (Song and Wessel, 2007). These were initially understood for their role in microRNA regulation, a process that uses the same molecular machinery as RNAi. MicroRNA regulation in sea urchin development has been studied extensively indicating that all the required proteins for building a RISC complex are present in the embryo (Song et al., 2012; Sampilo et al., 2018; Konrad et al., 2023).

Improved efficiency of RNAi reagents in a variety of organisms led to the commercial introduction of a double stranded RNA with a 2-base 3’ DNA nucleotide overhang on one strand (Kim et al., 2005; Amarzguioui et al., 2006). The availability of these Dicer-substrate interfering RNA (DsiRNA) products prompted us to investigate the possibility of using DsiRNAs for RNAi gene knockdown in the developing urchin embryo. DsiRNAs consist of a 25bp RNA-DNA hybrid sense strand where two dNTPs make up the last two bases of the 3’ end, and a 27bp RNA antisense strand that overhangs the 5’ end of the sense strand by two bases. The modification of the sense strand provides stability to the complex and helps bias against sense strand integration into RISC. The length ensures that processing and loading into the RISC complex includes Dicer (Kim et al., 2005; Amarzguioui et al., 2006).

We report successful adaptation of DsiRNAs for perturbation experiments with sea urchin embryos. A visual experiment reports the efficacy of DsiRNAs in initiating transcript degradation using an exogenously supplied GFP mRNA. With knowledge that the RNAi-based knockdown works in sea urchins, we used *pks1*, as an endogenous gene encoding an enzyme necessary for pigment production in the embryo, to identify and optimize the choice of sequences that are likely to work in the DsiRNA protocol. An effective DsiRNA was found for this gene indicating endogenous transcripts can be degraded in the developing embryo by RNAi. Methods were developed for monitoring the loss of a functional mRNA independent of the requirement for an observed phenotype.

We turned next to targeting two well-known signaling pathways for further tests on endogenous genes. Nodal and Lefty are well documented as essential for establishing of the dorsal-ventral axis in the sea urchin embryo (Duboc et al., 2004; Duboc et al., 2005; Range et al., 2007; Nam et al., 2007; Duboc et al., 2008; Bolouri and Davidson, 2009; Su et al., 2009; Duboc et al., 2010; Yaguchi et al., 2010; Molina et al., 2013; Materna et al., 2013; Coffman et al., 2014; Piacentino et al., 2015; Molina and Lepage, 2020; Floc’hlay et al., 2021; Pieplow et al., 2021).

Nodal establishes the ventral side of the ectoderm beginning early in cleavage. Molecular events following the expression of *nodal* include activation of *lefty* expression, and Lefty, in turn, antagonizes the function of the Nodal signal (Oulad-Abdelghani et al., 1998; Duboc et al., 2008), preventing Nodal from reaching the dorsal side. If Nodal is knocked down by a MASO or using an inhibitor, the embryo fails to establish the dorsal-ventral axis, and without any asymmetry the embryo adapts a radialized phenotype. If, on the other hand, Lefty is knocked down, Nodal function expands throughout and is said to radialize because now the embryo is entirely ventral. *Nodal* and *lefty* DsiRNA knockdowns match the known effects of perturbing expression of these genes using other perturbants with their well-known phenotypes. A dilution series showed that the DsiRNAs exhibit a graded inhibitory activity. To determine if a DsiRNA, after being injected at fertilization, can function later in development, the *chat* gene was targeted. This gene encodes an enzyme necessary for acetylcholine production and is not activated until after 18-20 hpf in *Lv* (Slota and McClay, 2018). DsiRNA directed to *chat* mRNA destroys that RNA indicating this approach can be used throughout development to the larval stage, at least, in the sea urchin embryo.

## Results

### DsiRNA-based knockdown is effective in sea urchin embryos

With knowledge that the sea urchin embryo expresses Dicer (Song and Wessel, 2007), and with commercial availability of DsiRNA oligomers, we tested an artificial system in the embryo. An mRNA encoding a transmembrane sequence fused to *gfp* (Saunders and McClay, 2014) was injected into the zygote and the membrane anchored-GFP fluorescently labeled all cell membranes (**Fig. 1).** At the 2-cell stage of these embryos, one blastomere was injected with a DsiRNA oligo targeting the *membrane*-*gfp* mRNA, and with Rhodamine isothocyanate-dextran (RITC-dextran) to fluorescently tag the DsiRNA-injected cell and its progeny. Embryos at 4, 6, 8 and 20 hpf were imaged to determine GFP fluorescence. As shown in **Fig. 1**, GFP expression in the DsiRNA-injected half-embryo is reduced to background over time indicating RNAi-mediated destruction of the *membrane-gfp* mRNA. And, since the added *membrane-gfp* mRNA was at a high concentration relative to endogenous transcript concentrations, the DsiRNA knockdown capability is substantial. The zygote had roughly 90 min. of time to accumulate membrane-GFP protein before adding the DsiRNA oligo, so at 4 hpf membrane GFP was expressed in all cells.

**Fig. 1.**
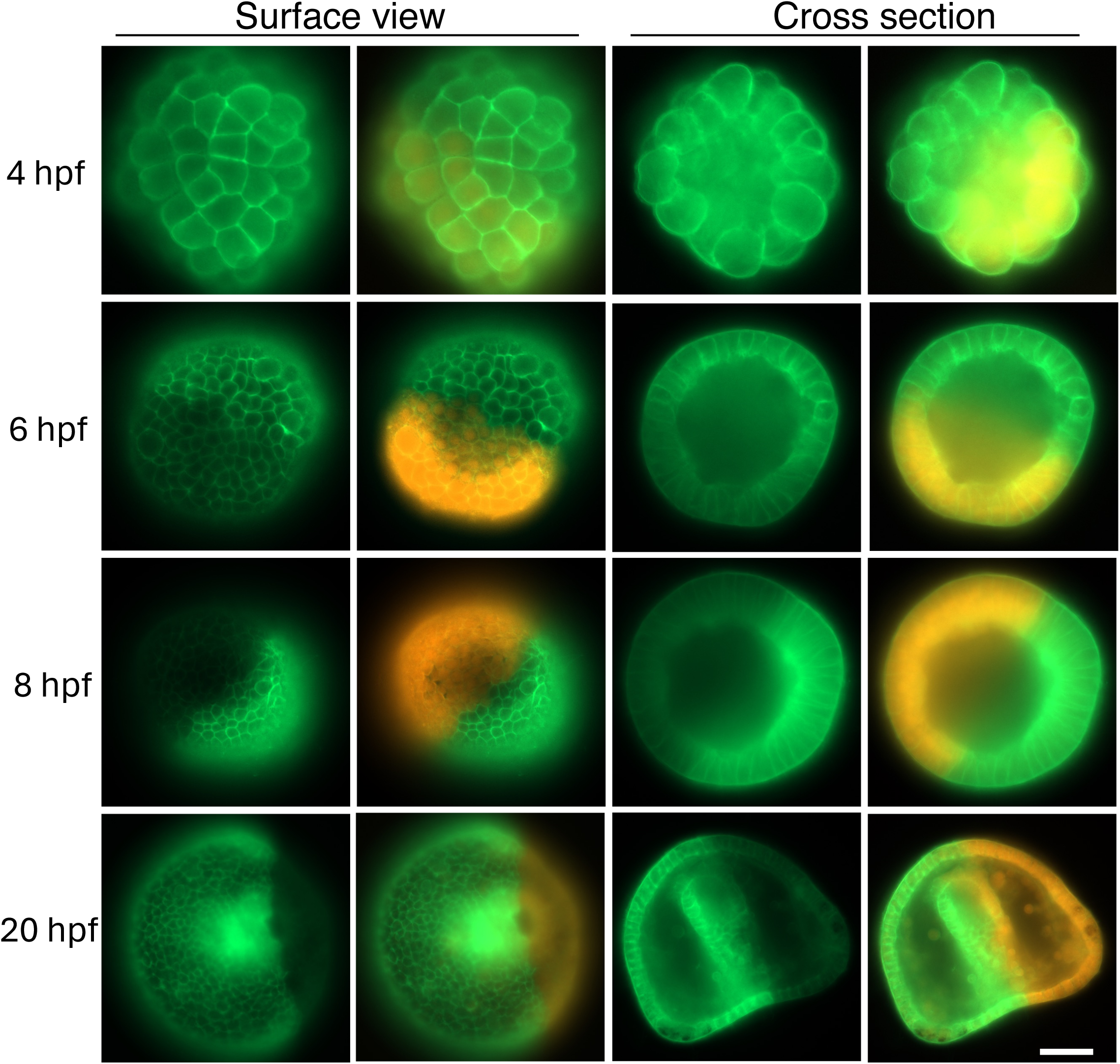
DsiRNA to GFP destroys a transmembrane sequence-GFP mRNA. Sea urchin eggs were injected with an mRNA expressing membrane-GFP. At the 2-cell stage, one of the two blastomeres received DsiRNA targeting *gfp* mRNA (450 ng/µl) plus RITC-dextran as a tracer. At 4, 6, 8, and 20 hpf embryos were imaged both with a surface view (minus or plus the tracer (orange)), and a mid saggital section of the embryo (minus and plus the tracer). As can be seen at 6 hpf there is a significant drop in GFP fluorescence (quantified in **Fig. S1** on six embryos per timepoint), a further drop at 8 hpf, and the fluorescent side at 20 hpf is in strong contrast to the DsiRNA injected side. Bar = 25 µm.

By 6 hpf, progeny of the injected blastomere exhibited significant loss of GFP fluorescence (**Fig. 1 and Fig. S1)**, and by 8 hpf almost all fluorescence was absent in the Dsi-RNA injected half, indicating that the destruction of the mRNA was essentially complete. The GFP protein is known to turn over relatively slowly in sea urchin embryos (a half-life of about 8.3 hrs in *Strongylocentrotus purpuratus* at 15°C (Arnone et al., 1997)), and the high level of introduced membrane-*gfp* mRNA in this artificial system makes it difficult to know how quickly the introduced DsiRNA destroys the mRNA. However, assuming the reported long half-life of the fluorescent GFP, the DsiRNA-destruction must have been significant well before 8 hpf indicating a high efficiency of the RISC complex in depleting even a high concentration of mRNA. We conclude that RNAi works in sea urchin embryos and can generate fully penetrant knockdowns of even highly abundant transcripts.

### Optimization of endogenous mRNA knockdown with DsiRNA

To understand details of the DsiRNA-induced knockdown of an endogenous gene we chose *pks1*, a gene encoding an enzyme in the mesodermal pigment production pathway (Calestani et al., 2003; Calestani and Rogers, 2010). Experiments with MASOs showed that knockdown of *pks1* translation results in albino embryos (Calestani et al., 2003). CRISPR knockout also results in albino embryos (Oulhen and Wessel, 2016). To ask if DsiRNAs also are effective in knockdown of expression, DsiRNAs were designed using the selection criteria described in the methods and were tested by injection and initially scored based on pigmentation of larvae. **Fig. 2A,B** shows that injection of the chosen DsiRNA leads to development of albino larvae. The loss of *pks1* mRNA is shown using conventional RNA *in situ* hybridization with a digoxygenin probe. *pks1* is expressed in control embryos, but absent in DsiRNA-injected embryos (**Fig. 2C,D).** In four separate experiments 100% of injected embryos were devoid or almost devoid of *pks1* signal as seen by *in situ* hybridization. Occasionally, a few cells in embryos that likely received a lower injected volume of 75µM DsiRNA (estimated by FITC fluorescence intensity of the tracking dye) were seen expressing a very low level of *Pks1* (**Fig. S2),** indicating incomplete knockdown. Even then, however, the level of knockdown is almost complete. This outcome is reinforced by the phenotype of larvae. The population of DsiRNA-treated larvae in each of the four experiments are albino with only a few of hundreds of larvae showing one or two slightly pigmented cells.

**Fig. 2.**
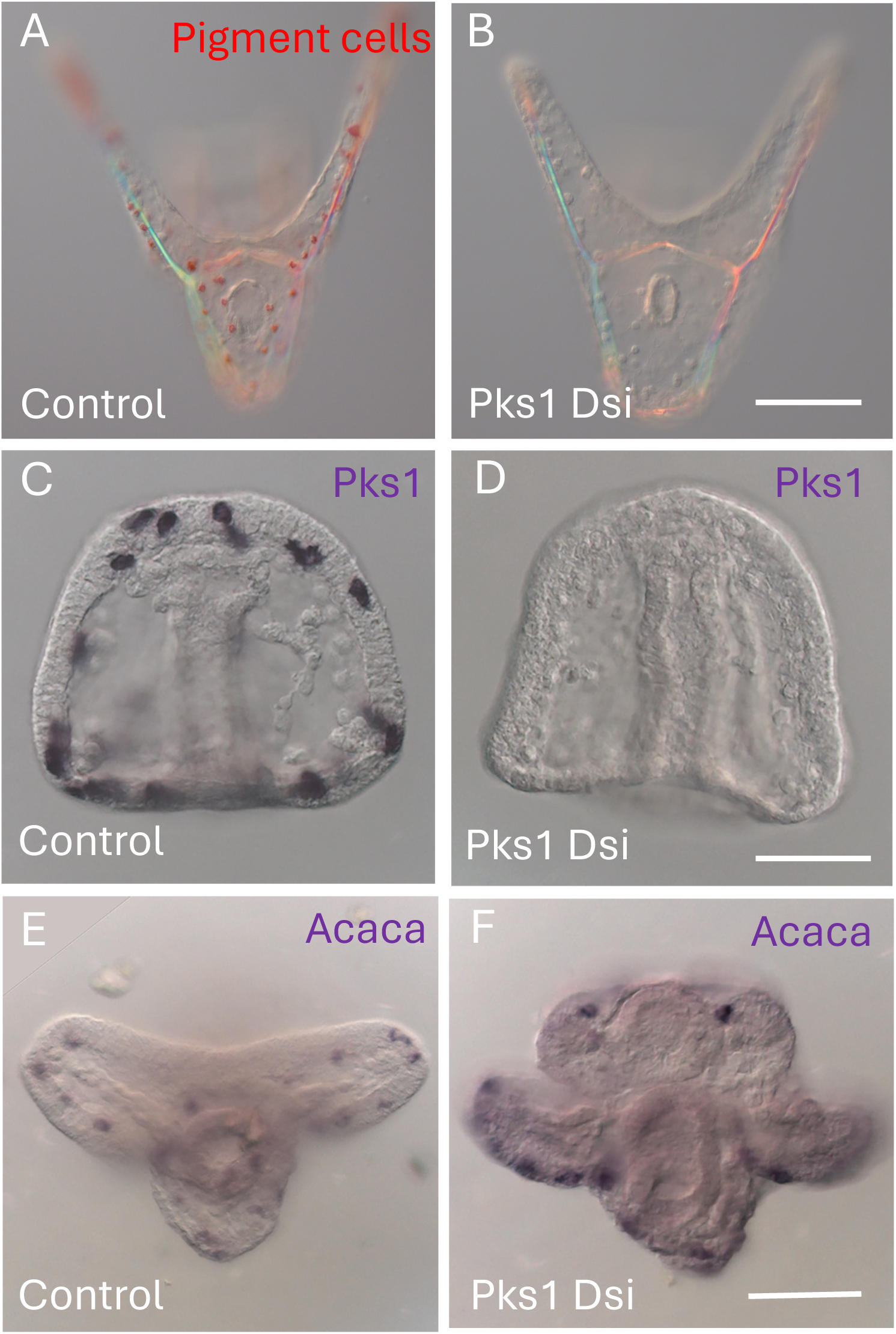
DsiRNA to *pks1* strongly reduces expression of the mRNA for this enzyme. (A,B) 48 hpf larvae with pigment cells in red in the control **(A),** and absent in the larva injected with DsiRNA to *pks1 (n* imaged with pigment cells: 0/12) **(B). (C,D)** In situ of *pks1* at 20 hpf showing presence of mRNA to *pks1* in the control **(C),** but absent from the embryo injected with DsiRNA to *pks1 (n =* 0/12 had *pks* expression) **(D). (E,F)** In situ of *acaca,* a gene expressed specifically in pigment cells (**Fig. S2),** showing its presence in both control and in embryos injected with DsiRNA to *pks1.* (n=12/12 embryos imaged). Thus, the Dsi removes the mRNA to *pks1*, but does not eliminate presumptive pigment cells.

A qPCR protocol was also adapted for the DsiRNA experiments (see methods). It too showed a greater than 90% drop in the amount of *pks1* mRNA present in the knockdown embryos relative to controls (**Fig. S3A)**. Results of verification of pks1 knockdown by qPCR are consistent with information we gleaned from *in situ* hybridization. Although qPCR provides a quantitative verification of a knockdown, we prefer *in situ* hybridization as a method of verification owing to low input and the added value of spatial and morphological information in developmental contexts. The recent addition of Hybridization Chain Reaction (HCR) *in situ* analysis also was analyzed for its possible efficacy in monitoring DsiRNA knockdowns. **Fig. S2C,D** shows the result of that analysis. We were surprised to discover that in DsiRNA-injected embryos (injected in the same experiment that were analyzed with conventional Digoxygenin-probes), there was a fluorescent signal indicating the presence of *pks1* in the DsiRNA-injected embryos. We then realized there is a difference in the probes between Digoxygenin-based and HCR-based *in situs*. The Digoxygenin probes require targeted RNA to be greater than 500 bp in length while the HCR probes are designed to amplify recognition of many short oligonucleotide sequences along the mRNA (Molecular Instruments, Inc.). For Digoxygenin-based *in situs* to work the mRNA must be intact, while HCR probes can detect short RNA fragments of the DsiRNA-degraded RNA. Thus, while the *pks* DsiRNA knockdowns had the phenotype, and the knockdown was confirmed by standard Digoxygenin *in situ* analysis, and by qPCR, HCR-probed DsiRNA knockdowns showed the presence of *pks* fragments, a signal that is like the HCR signal of control embryos (**Fig. S2).**

Embryos were further tested with a probe to *acaca*, (acetyl CoA-carboxylase), a gene that, like *pks1,* is expressed exclusively in pigment cells (**Fig. S2E,F).** Knockdown of *pks1* has no effect on expression of *acaca* indicating that pigment cells are specified in the knockdowns, but these cells are specifically devoid of intact *pks1*.

### Nodal and Lefty DsiRNA knockdowns phenocopy knockdowns using MASOs or drug inhibitors

A well-understood signaling system in the sea urchin embryo is Nodal and Lefty signaling. The Lepage group identified Nodal as the crucial signal that establishes dorsal-ventral and left-right axes in sea urchin development (Duboc et al., 2004; Duboc et al., 2005; Duboc et al., 2008; Duboc et al., 2010; Floc’hlay et al., 2021). *Nodal* is expressed by ventral ectoderm and establishes a “community effect” signaling center during cleavage (Gurdon, 1988; Bolouri and Davidson, 2009), and that Nodal signaling initiates specification of ventral tissues. Among the genes activated downstream of the Nodal signal, *lefty,* an anti-Nodal signal, restricts the reach of Nodal to the ventral ectoderm, ventral mesoderm and ventral endoderm (Duboc et al., 2004; Duboc et al., 2008; Duboc et al., 2010). Inhibition of *nodal* translation with a MASO or a specific drug inhibitor radializes the embryo because ventral genes fail to be activated.

Inhibition of *lefty* translation with a MASO allows the Nodal signal to expand its reach into the presumptive dorsal territory and activate ventral-specific genes in cells that would otherwise be dorsal, resulting in a radialized embryo that is entirely ventral in identity. Dorsal identity is established by expression of BMP2/4 that oddly enough, is expressed downstream of Nodal in the ventral ectoderm. BMP2/4 protein is transported to the dorsal half of the embryo, possibly using Chordin as a chaperone (Madamanchi et al., 2021), and likely employing Glypican5 to facilitate that transport (Lapraz, 2009). The BMP2/4 signal is activated in the dorsal half via a tolloid-like protease that removes Chordin from the BMP2/4 ligand which then establishes the dorsal GRN (Duboc et al., 2004; Duboc et al., 2008; Duboc et al., 2010).

**Figs. 3A and 3B** illustrate the phenotypic outcome to mesoderm specification following injection of a dilution series of MASOs to *nodal* or *lefty*. The experiments score the expression of two mesodermal genes. *Irf4* is a transcription factor expressed by ventrally specified blastocoelar cells (**Fig. S2G)** (Hibino et al., 2006; Allen et al., 2022), with the Nodal signal required for its expression **(Fig. 3A).** Pigment cells expressing *pks1* arise from dorsal mesoderm. Inhibition of Lefty results in excess Nodal reaching the presumptive dorsal mesoderm, and that leads to an inhibition of *pks1* expression **(Fig 3B)**. As can be seen in **Fig. 3B**, a 1:36 dilution of MASO still results in insufficient Lefty protein needed to block Nodal diffusion to the dorsal mesoderm. Dilution of the *nodal* MASO shows that ventral mesoderm specification is sensitive to Nodal concentration, i.e. the number of blastocoelar cells increases from total absence to control levels over about a 10-fold dilution range of the MASO. The *nodal* inhibition is more complicated than the Lefty knockdown since Nodal also controls the expression of *lefty* and *BMP2/4.* Excess BMP2/4, as shown by the Lepage group, results in a dorsal radialized embryo **(Fig. 3C)** (Duboc et al., 2004; Duboc et al., 2008; Lapraz, 2009; Duboc et al., 2010).

**Fig. 3.**
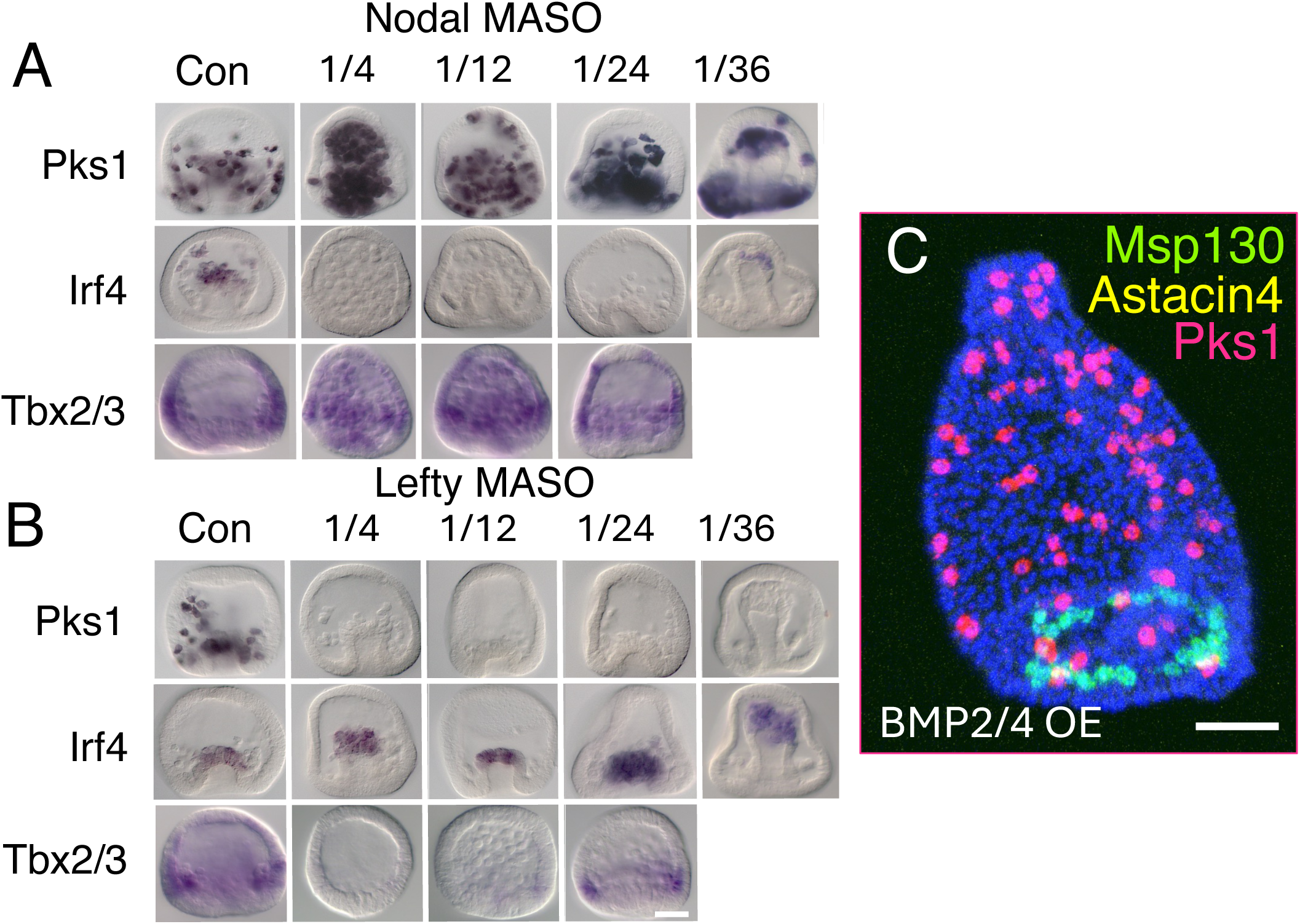
Nodal, Lefty, and BMP signaling establish dorsal-ventral axis. (A) A MASO to *nodal* was injected into zygotes in a dilution series beginning with a concentration of 0.75mM. At 20 hpf embryos were fixed and in in situs performed with a dorsal mesoderm marker (*pks1*), ventral mesoderm marker (*irf4*), and a dorsal ectoderm marker (*tbx2/3*) (Gross et al., 2003). Over the entire dilution range of the *nodal* MASO, excess dorsal pigment cells are specified, and no ventral mesoderm cells are specified until the 1:36 dilution of the MASO. The dorsal marker is present throughout. **(B)** A MASO to *lefty* was injected into zygotes in a dilution series beginning with a concentration of 0.75mM. Normally, Lefty knockdown results in ventralization of the embryo because too much Nodal reaches the otherwise dorsal half of the mesoderm resulting in absence of pigment cells (*pks1* marker). Even at a 1/36 dilution of the Lefty MASO there is insufficient Lefty protein to block this ventralization. The ventral marker is present throughout as expected. The ectoderm is less sensitive to excess Nodal in that *tbx2/3* returns at a Lefty MASO dilution of 1/24. (**C**). Overexpression of BMP2/4 results in dorsal radialization of the embryo producing pigment cells around the entire embryo (red), a radial arrangement of skeletogenic cells (green) and an absence of a ventral mesoderm marker (yellow – Astacin4 is not expressed). At each dilution all embryos imaged were similar in expression of a selected marker (≥ 6 images per dilution). Line=25µm.

With the above as a control series in *Lv*, we designed and tested three DsiRNA candidates each for blocking expression of *nodal* or *lefty*. One DsiRNA candidate targeting each gene was most effective in matching the MASO outcomes (**Fig. 4)**. Each of the six DsiRNAs were also tested by *in situ* to determine whether there was visual evidence of mRNA knockdown of *nodal* or *lefty.* **Figs. 4D and H** shows that the DsiRNAs that gave a radialized phenotype for each (*nodal* DsiRNA-3, and *lefty* DsiRNA-6) were devoid of intact mRNA as seen by *in situ* hybridization while the other DsiRNA candidates failed to produce a radialized phenotype and were positive for *nodal* or *lefty* expression by *in situ*. *Lefty* DsiRNA-4 gave a weak phenotype and its expression by *in situ* was reduced relative to controls. **Fig. S3A** shows the qPCR analysis of the *lefty* and *nodal* DsiRNAs that produced phenotypes. Again, there was a greater than 90% knockdown of both. One of the repeats of the *lefty* knockdown was very different from the other two accounting for the large variation in that analysis. To further analyze knockdowns, expression of *pks1* in *nodal* or *lefty* DsiRNA knockdowns was tested by qPCR (**Fig. S3B**). As expected, in the *lefty* knockdown there was a significant loss of *pks1* while in a *nodal* knockdown there was excess *pks1* expression (**Fig. S3B). Fig. S4** shows the phenotypic outcome of the *nodal* and *lefty* DsiRNA inhibition as seen by HCR. Again, because the HCR probes recognize RNA fragments, both the control and the DsiRNA knockdowns show the presence of *lefty* and *nodal*, further indicating that as a control, qPCR and conventional Digoxygenin *in situ* probes reveal the knockdown status but HCR *in situ* analysis does not.

**Fig. 4.**
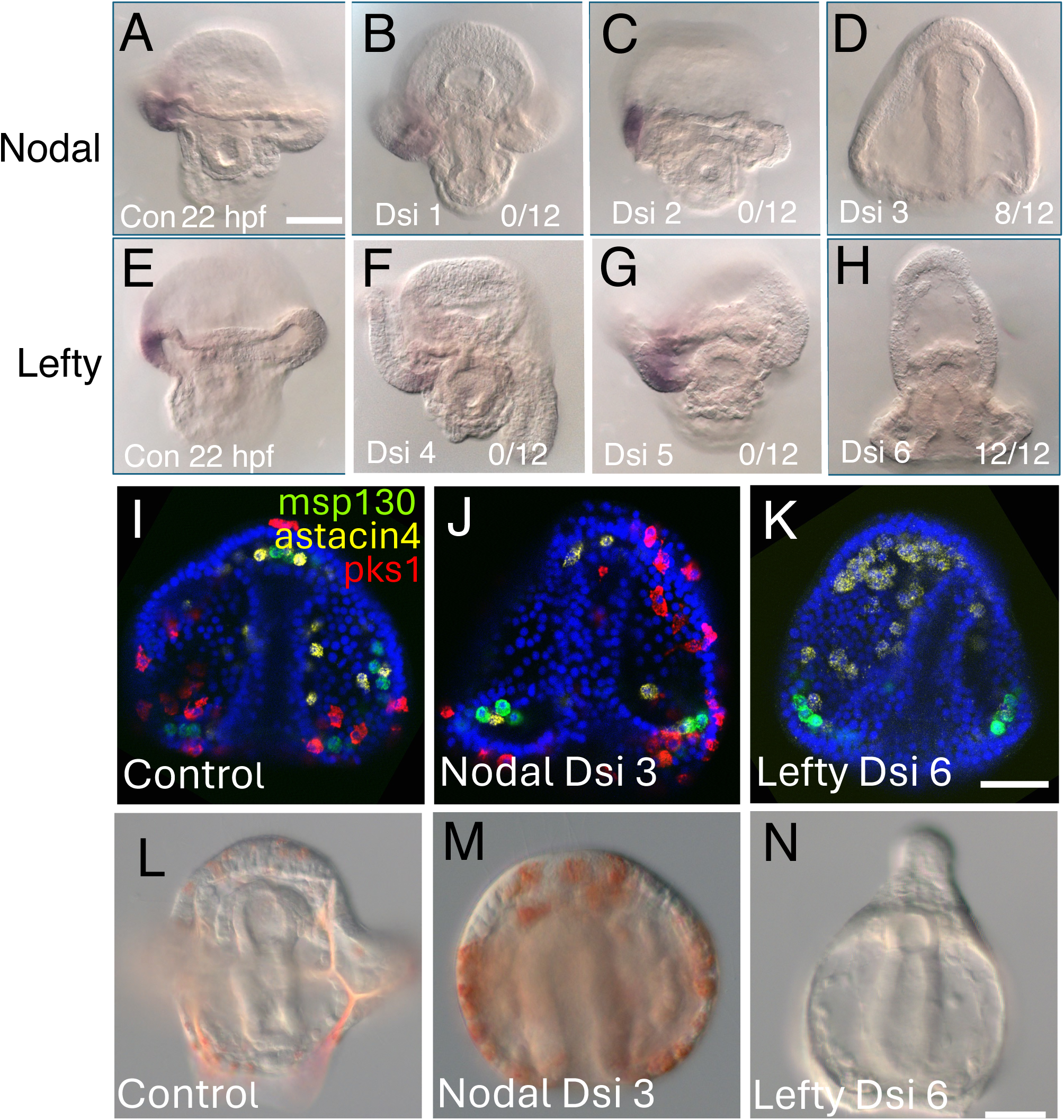
**DsiRNA to *nodal* and *lefty*: Functional DsiRNA destroys mRNA. (A**) Control Digoxygenin *in situ* showing *nodal* expression on the right side. (**B-D)** DsiRNA candidates designed for *nodal.* **B** and **C** failed to radalize the embryos and showed *nodal* expression reflecting failure as a candidate. Numbers indicate embryos sampled that give a radialized phenotype. The embryo in **D** is radialized, and there is no visible expression of *nodal* by *in situ* confirming that DsiRNA 3 functions to destroy *nodal* mRNA. Further experiments used DsiRNA 3 only (n = ≥6 embryos imaged per 100 injected embryos per experiment of four experimental repeats and knockdown results by in situ were consistent. (**E**) Control expression of *lefty* on the right side seen by Digoxygenin *in situ* **(F-H)** *lefty in situs* of embryos injected with *lefty* DsiRNA candidates. **F** and **G** show candidates that fail to knock down *lefty* (0/12) while DsiRNA 6 (12/12) in **H** shows absence of *lefty* mRNA, and the embryo is radialized. Further experiments used DsiRNA 6 (n ≥ 6 embryos imaged in four separate experiments with more than 100 injected embryos subjected to *in situ* in each experiment and results of each experiment were consistent with radialization and a *lefty* knockdown. (**I**) A control embryo probed by HCR for dorsal *pks1* (red), ventral *astacin4* (yellow) and skeletogenic cells (*msp130,* green). (**J**) DsiRNA 3 to *nodal* shows an over expression of dorsal pigment cells and reduction of ventral blastocoelar cells. (**K**) DsiRNA 6 to *lefty* shows an absence of dorsal pigment cells and an overabundance of ventral blastocoelar cells. (**L-N).** Brightfield images of control (**L**), *nodal* DsiRNA knockdown (radialized and too many pigment cells) (**M**), and *lefty* DsiRNA knockdown (strongly radialized and no pigment cells) (**N**). (n ≥ 7 embryos imaged with each treatment). Bars = 25 µm.

We next used HCR analysis in a different way; to image the patterning outcome of the nodal*-lefty* experiment. In **Figs. 4I-K** each embryo is stained by HCR probes to a dorsal mesoderm marker (*pks1*, red), ventral mesoderm marker (*astacin4*, yellow), and a skeletogenic cell marker (*msp130*, green). For each marker, lineage specific expression is shown in scRNA-seq profiles **(Fig. S2, S4)**. *Pks1,* the dorsal marker, is missing in the *lefty* DsiRNA knockdown, and the ventral marker (*astacin4*) is greatly reduced in the *nodal* DsiRNA knockdown. In **Fig. 4L-N**, the radial phenotypes are seen in DsiRNA knockdowns of *lefty* and *nodal* and the *nodal* knockdown has extra pigment cells while the *lefty* knockdown is albino.

The *nodal* and *lefty* DsiRNAs that gave phenotypes and eliminated mRNAs were further studied in a dilution series (**Fig. 5**). For *lefty*, a concentration of DsiRNA four-fold higher than the concentration necessary to eliminate *pks1* expression showed embryos that lacked any appearance of toxic effects. For both *nodal* and *lefty*, a concentration of 75-100 µM of the DsiRNA was effective in producing the known phenotype. Finally, for both, the dilution series showed that the knockdown was graded and concentration dependent. The territory-specific HCR probes were used to quantify the dorsal-ventral signaling outcomes of three mesoderm cell types in the same embryo. We counted the number of dorsal pigment cells (red), ventral blastocoelar cells (yellow) and skeletogenic cells (green) in the confocal-stacked images at each dilution. The number of skeletogenic cells remains at a similar level throughout indicating that their number is unaffected by *nodal* or *lefty* concentration changes. In the DsiRNA dilution series of *lefty*, an absence of pigment cells is observed down to an injected concentration of 100 µM, with further dilution of the DsiRNA producing a graded response. Those same embryos, when blastocoelar cells were scored, had an excess number of ventral blastocoelar cells at higher concentrations of *lefty* DsiRNA, and dilutions produced a graded response. The knockdown was dose dependent over a 20-fold range of concentrations. The *Nodal* DsiRNA dilution series also displayed graded response, though less dramatic than the *Lefty* knockdown. These dilution series confirm and expand the largely qualitative analysis of the Lepage group (Floc’hlay et al., 2021) by showing how quantitative changes in *nodal* and *lefty* impact dorsal and ventral mesoderm patterning over a wide range of Nodal or Lefty concentrations.

**Fig. 5.**
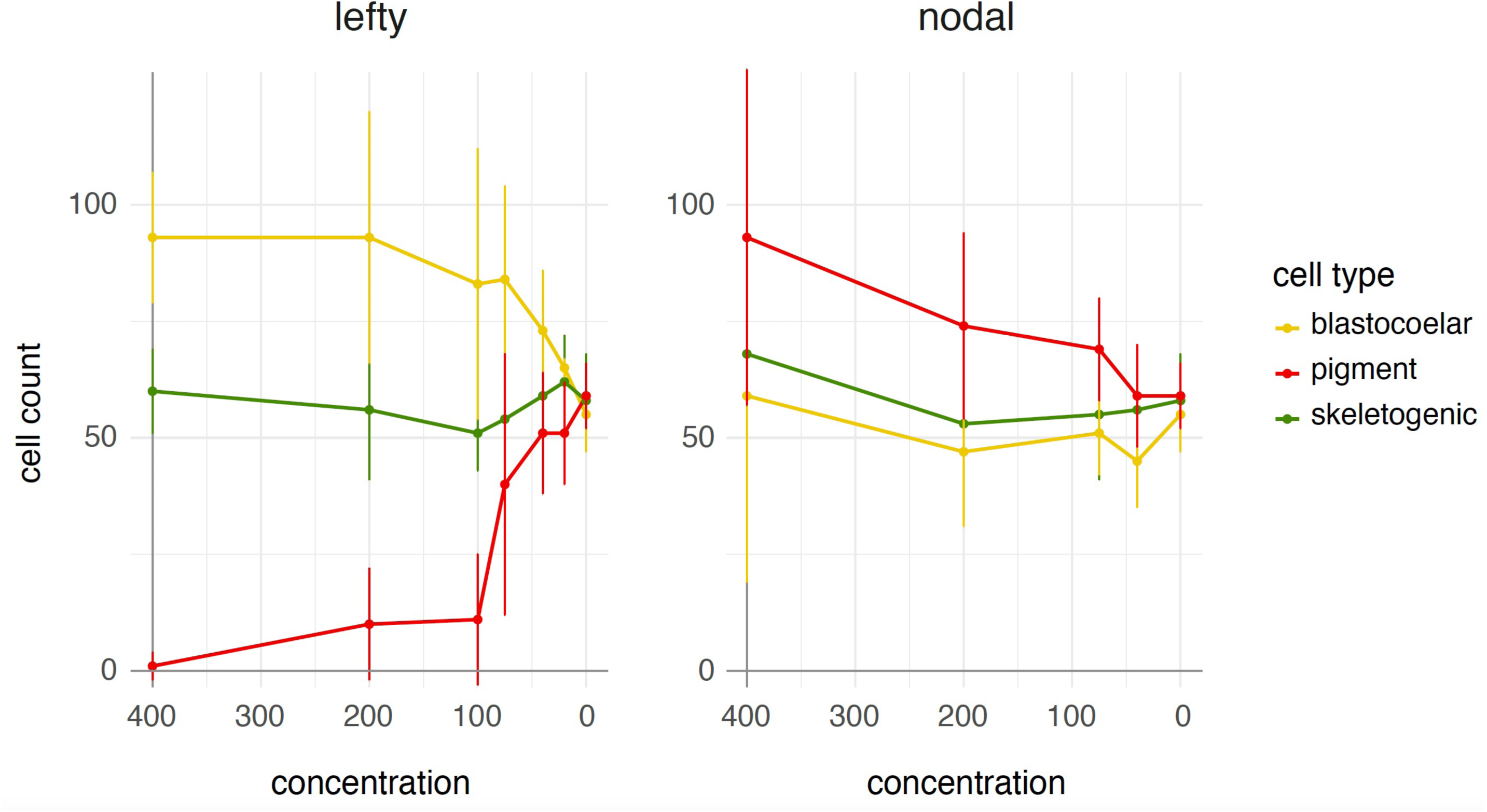
A quantitative assessment of *nodal* and *lefty* knockdowns over a dilution range of DsiRNA. Blastocoelar, pigment and skeletogenic cells were quantified by HCR in embryos injected over a 20-fold dilution range of *nodal* and *lefty* DsiRNA beginning at 400 µM. Cells were counted from confocal image stacks (7 embryos per concentration) of DsiRNA with embryos fixed at 20 hpf (hours post fertilization). Pigment cells (dorsal) are red, blastocoelar cells (ventral) are yellow, and skeletogenic cells (included as a test of possible toxic effects) are green. Over the entire concentration range, including a concentration 4-fold higher than necessary to eliminate all pigment cells, the skeletogenic cell number remained at a similar level. Both pigment cells and blastocoelar cell numbers were sensitive to DsiRNA concentrations. Each data point reflects the mean ± std. dev. of pigment, blastocoelar or skeletogenic cells in 7 embryos at each concentration.

### DsiRNA – RISC complexes operate from fertilization to at least the larval stage

Experiments so far indicate that DsiRNA works for mRNAs that are present early in development and it efficiently destroys mRNA. The *membrane-gfp* mRNA experiment introduced on the order of 10^5^ molecules of mRNA, a concentration that is at least 10-fold higher than abundant endogenous mRNAs (actin as an example (Steuerwald et al., 2000)). By 6 hpf most membrane-GFP is no longer present, and since GFP protein is relatively long-lived (8.3 hrs half-life in *S. purpuratus*, (Arnone et al., 1997)), the DsiRNA likely operated efficiently from the time of injection at fertilization.

To examine the ability of DsiRNA to operate later in development, we turned to a gene expressed late in development. *Chat* (choline O-acetyltransferase), an enzyme in the acetylcholine synthesis pathway, is first expressed in cholinergic neurons with the earliest detection occurring at about 18 hpf in *Lv* (Slota and McClay, 2018). A DsiRNA was designed to destroy *chat* mRNA and by conventional *in situ* analysis, embryos injected with the *chat-*directed DsiRNA, showed that *chat* is absent while control embryos display a robust expression of *chat* at 24 hpf, allowing plenty of time for *chat* to accumulate in both experimentals and controls (**Fig. S5**). We conclude that DsiRNA produces knockdowns as far as 24hpf into development and suspect that they may retain efficacy well beyond that point, as they are reported to retain activity for several days in mammalian cell cultures (Ambardekar et al., 2018).

## Discussion

DsiRNAs trigger destruction of targeted transcripts in the sea urchin. That is shown with data from the visual GFP construct, knockdown of *pks1*, an enzyme necessary for production of pigmented cells, knockdowns of *nodal* and *lefty* affecting acquisition of the dorsal-ventral axis, and knockdown of *chat*, an enzyme expressed late in development. The DsiRNA results phenocopy known outcomes from other knockdown methodologies. A dilution series of the DsiRNAs to *nodal* and *lefty* indicated that the outcome is a quantitative measure of dorsal or ventral signaling perturbations in that the number of pigment cells or blastocoelar cells produced reflected the level of inhibition. Toxic effects were absent or undetected over a 20-fold range in DsiRNA concentrations, including a 4-fold higher concentration than necessary to eliminate production of pigment cells. There were no observed developmental delays or abnormal effects on skeletogenic cell number.

Two major experimental advantages of DsiRNA reagents over MASOs are the way the inhibition is achieved and the ease of use. MASOs block translation. DsiRNA reagents, by contrast, destroy the transcript. This makes it possible to validate knockdown by a simple *in situ* analysis. While Digoxygenin *in situ* hybridization analysis is only semi-quantitative, critical information is supplied using that tool as to whether the mRNA in question is significantly reduced, and in the expected location in the embryo. The completeness of a MASO translation knockdown is, by contrast, usually unknown. Only if an antibody to the targeted protein is available, or if the MASO is a splice blocking reagent, can the investigator ascertain the relative completion of MASO knockdown. Most MASO experiments rely on two MASOs with the logic that if both produce the same, or a similar phenotype, the likelihood of off-target effect explaining that phenotype is small. Usually, one of the two MASOs is more potent, and the assumption is made that it blocks translation more completely than the other, but in neither case are those assumptions validated. With RNAi-mediated knockdowns, the investigator can easily test for on-target knockdown, providing more confidence in the results.

Of the three tests employed to determine destruction of the targeted mRNA, each has utility. We find digoxygenin *in situ* analysis to be the simplest and most informative because it records the presence or absence, or leak-through expression of the targeted mRNA, and it provides spatial expression information. qPCR also is effective but can be cumbersome. We used an approach that consisted of mRNA isolation by pulldown with magnetic poly-dT beads ahead of reverse transcription, paired with primers targeting a region of the mRNA 5’ of the DsiRNA sequence. This allows a quantitative comparison between full length mRNA in the controls, and fragmented mRNA in the DsiRNA embryos. qPCR on total RNA often gave confusing results. For example, in the *lefty* knockdown an excess amount of *nodal* mRNA is produced, and in excess Nodal, there is an increased production of *lefty* mRNA. Using traditional qPCR approaches, this created the appearance of a dramatic increase in *lefty* expression in the *lefty* knockdown, likely owing to amplification of fragmented mRNA. The cumbersome aspect of the qPCR approach is the necessity of starting with about 200 injected embryos to obtain enough mRNA for the analysis. With Digoxygenin *in situ* analysis, we obtain the information needed with about 10 injected embryos. HCR analysis, which uses multiple probes that recognize short sequences along the mRNA (Molecular Instruments, Inc.) is not useful as a measure of efficacy in RNAi-mediated knockdown. However, if Digoxygenin *in situ* analysis shows an absence of a signal while HCR shows presence, this is strong evidence that the target mRNA has indeed been fragmented. Other approaches can likely also work for validating a knockdown, such as probe-based qPCR with relatively long amplicons, or RNAseq (depending on the specific technology and library preparation). Regardless of the technology, we suggest considering ahead of time whether fragmented target RNAs are likely to be distinguishable from intact RNAs in the data.

The ability of DsiRNAs to mediate knockdown in the sea urchin embryo appears not to be limited by target transcript abundance: we calculate that about 4.8 x 10^5^ copies of *membrane-gfp* mRNA were delivered into each zygote in the GFP knockdown experiments. Estimates of the total number of mRNAs in a cell vary, but a quantitative measure in mice gave 4-8 x 10^5^ total mRNA copies per cell (Carter et al., 2005; Milo, 2015). Estimates of the number of β-actin mRNAs in mouse and human metaphase II oocytes range from 1.2 x 10^3^ to 3.8 x 10^4^ copies (Steuerwald et al., 2000). The injected *membrane-gfp* mRNA therefore almost certainly exceeds the abundance of all endogenous mRNAs in the zygote by a significant margin. The zygotes have about 90 minutes of protein synthesis before the GFP-targeted DsiRNA is injected into one blastomere at the 2-cell stage, providing the first opportunity for RNAi-mediated decay. The control side of the embryo continually produces membrane-GFP protein, exhibiting a tenfold brighter fluorescence signal at 20 hpf than the control side at 4 hpf. In the progeny of the DsiRNA-injected cell, the fluorescent signal is significantly reduced by 6 hpf, and essentially gone by 8 hpf. Given that GFP has a reported half-life of 8.3 hours in the developing embryo (Arnone et al., 1997), the simplest explanation for these results is a rapid and early destruction of the introduced mRNA. Similarly, the knockdown of *nodal*, whose earliest transcripts are detectable at 32-cell stage, indicates that DsiRNAs effectively mediate knockdown very early in development. The knockdown of *chat* indicates that the reagent is sufficiently stable to destroy an mRNA that isn’t expressed until at least 18 hpf. Thus, DsiRNA-mediated knockdown enables perturbation of targets present very early and very late in the developing sea urchin and is robust enough to destroy even transcripts whose abundance vastly exceeds the abundance of endogenous mRNAs.

Our design process for generating DsiRNAs has been and is continually evolving, and our current selection protocol (see Materials and Methods) should be viewed in that light. We arrived at the current approach through some trial and error, with reasoning behind each step provided below. Our initial approach involved two major steps. First, we used the DsiRNA design tool provided by Integrated DNA technologies (https://www.idtdna.com/site/order/tool/index/DSIRNA_CUSTOM) to generate candidate sequences against the full cDNA sequence of a given target gene. This tool produces a maximum of 50 possible DsiRNA designs per input sequence. Secondly, we used NCBI’s BLAST tool (https://blast.ncbi.nlm.nih.gov/Blast.cgi) to check each candidate for possible off-target hits against both the genomic and non-redundant databases for *Lv* (see Materials and Methods for details). This step is particularly important in sea urchins, whose genomes exhibit a high level (about 1 in 20 bases) of polymorphism (Britten et al., 1978; Sodergren et al., 2006).

This initial process resulted in multiple functional DsiRNA designs, but with only ∼1 out of 3 designs producing a phenotype for each gene tested. As the algorithm used in the IDT DsiRNA design tool is proprietary, we began checking designs against the more transparent shRNA design tool found on the Genomic Perturbation Platform provided by the RNAi consortium at the Broad Institute (GPP, available at https://portals.broadinstitute.org/gpp/public/seq/search) in hopes of generating a higher success rate by adding that tool’s known parameters. The algorithm employed in this tool is designed to score 21-mers while DsiRNAs are 27-mers. As a result, output from the tool provides multiple intrinsic activity scores to evaluate for each design entered (see Materials and Methods). We also removed UTRs from consideration due to a higher rate of polymorphism, using only the CDS from each gene in DsiRNA design. Incorporating all these elements together appears to provide a higher rate of successful design relative to our initial attempts. DsiRNAs against more than 20 additional genes in ongoing projects in the lab have a success rate topping 50%. It is important to note that even with all these design elements considered, there is no guarantee of each reagent producing a highly penetrant knockdown for the gene in question. We still design multiple DsiRNAs for each targeted gene. And importantly, conventional in situ analysis is a most useful tool to help decide whether to go forward with a DsiRNA in a project.

In using DsiRNA, as with any experimental reagent, off-target effects may be present. The genome blast step in choosing a DsiRNA helps in that regard because it warns us to avoid sequences that may recognize unwanted genes. In singlicate gene perturbation where the function of a gene is well-studied, as are the genes perturbed here, a knockout that phenocopies knockouts produced by other methods provides near certainty that the observed phenotype is due to an on-target effect. This changes with the targeting of genes for which a knockout phenotype is not clearly defined. Thankfully, RNAi is a mature technology in other systems, and best practices are well-established. Accordingly, for genes whose knockdown phenotype is not known, there are several ways of validating on-target performance. A common phenotype between multiple validated DsiRNAs is strong evidence that the on-target knockdown is responsible for the observed phenotype. Validation via orthogonal methods, wherein two distinct technologies are used to target the same gene, also provides strong support for causally linking a perturbed gene and resulting phenotype.

Both with MASOs and with DsiRNAs, a dilution series of *lefty* and *nodal* perturbations indicate a graded effect on signaling. As pointed out in many papers from the Lepage lab, but especially in the (Floc’hlay et al., 2021) paper, additional molecules beyond Nodal and Lefty contribute to establishing the dorsal and ventral mesodermal territories. The intriguing feature is that the territorial boundaries arrive at a sharp cutoff despite the demonstration that there is a graded dilution effect. Other data over the years in our lab suggests that the location of that boundary, while sharp when established (based on *in situ* patterns displayed by either dorsal or ventral markers), may not be absolutely at the same location in all batches of embryos. In different batches of embryos examined at an equivalent timepoint we see variable numbers of pigment and blastocoelar cells exhibited by one batch relative to another, suggesting the possibility that the precise location of that territorial boundary may differ a bit from one genetic background to another, but within a batch most embryos quantitatively demonstrate a similar number of pigment cells and blastocoelar cells. In that sense, the site of the boundary may be due to a threshold effect in which the exact site where Nodal no longer is above a threshold necessary for its transduction may vary in different genetic backgrounds.

MASOs have been a great resource for the sea urchin community, as without them the embryonic GRN would be very incomplete relative to its current status. Our lab has used well over 100 MASOs in our contributions to that systems analysis over the years. At the same time, we have always been concerned with the inherent lack of knowledge of the completeness of the translation block. Additionally, Gene Tools, Inc. warns investigators that the higher the G content of a MASO, the more likely the MASO will precipitate (Gene Tools, Inc), causing stock solution concentrations to shift over time and causing injection needles to clog. We’ve observed that this occasionally makes results obtained with these reagents difficult to reproduce. Compared to MASOs, we find that DsiRNAs are significantly cheaper, less toxic, and more stable in storage. Further, it is not a new technology. As cited above, many studies have worked toward optimization of RNAi including use of DsiRNA. Beyond those attributes, however, is the value of having multiple ways of perturbing specific molecules in a biological system. The addition of DsiRNA adds a new tool that is simple to use, capable of targeting transcripts early and late in development, and has great promise for an embryonic system that relies on reverse genetics for molecular information on how development works.

## Materials and Methods

### Animals and embryos

Adult *Lytechinus variagatus* (*Lv)* animals were obtained either from the Duke Marine Laboratory, Beaufort, NC or from Reeftopia.Com in Florida. Gametes were obtained by injection of 0.5M KCl. Fertilization and culture of embryos was in artificial sea water (ASW) at 23°C.

### Selection of DsiRNA sequences

All DsiRNAs used in this study were obtained from Integrated DNA Technologies (IDT, www.idtdna.com). Candidate DsiRNAs are generated by entering the CDS for the gene of choice into the IDT DsiRNA design tool to obtain initial candidates for the gene in question (https://www.idtdna.com/site/order/tool/index/DSIRNA_CUSTOM). BLAST is used to check each candidate for any possible off target matches at both the genomic and transcript level (https://blast.ncbi.nlm.nih.gov/Blast.cgi). Candidates with E-values for any off-target hits >= 1.0, with perfect off-target alignments to another transcript >= 14 bases, and candidates with any alignment of >= 15 bases with <= 2 mismatches to an off-target transcript are eliminated from further consideration. Candidates that fall within a known polymorphic sequence are then eliminated. Candidate DsiRNA designs are then entered into the Broad Institute’s GPP shRNA picking tool to retrieve intrinsic activity scores for each 27-mer, using the reverse complement of the antisense strand sequence as the input. Special emphasis is placed on high intrinsic activity scores at positions 1, 5, and 3 of the candidate sequence. From these screened acceptable candidates, multiple DsiRNA designs per gene complementary to distinct, non-overlapping regions of the transcript are ordered and tested for specific knockdown activity.

### Sequences of DsiRNA and MASOs used in this study

The sequences used in this study are:

Membrane-GFP DsiRNA

rArArG rCrUrG rArCrC rCrUrG rArArG rUrUrC rArUrC rUrGCA

rUrGrC rArGrA rUrGrA rArCrU rUrCrA rGrGrG rUrCrA rGrCrU rUrGrC

LvPks1 DsiRNA

rCrUrU rCrUrU rGrGrA rUrUrC rArCrU rGrArC rUrArC rArACA

rUrGrU rUrGrU rArGrU rCrArG rUrGrA rArUrC rCrArA rGrArA rGrUrA

LvNodal DsiRNA

rArGrArArGrGrArGrArArGrGrArArCrArArCrGrCrArArACA

rUrGrUrUrUrGrCrGrUrUrGrUrUrCrCrUrUrCrUrCrCrUrUrCrUrUrG

LvLefty DsiRNA

rGrArCrUrUrArCrArArArUrGrGrArUrArUrUrCrUrUrGrATA

rUrArUrCrArArGrArArUrArUrCrCrArUrUrUrGrUrArArGrUrCrArA

LvChat DsiRNA

rGrGrUrUrGrArGrCrArUrCrUrArCrUrGrArArArUrArCrATT

rArArUrGrUrArUrUrUrCrArGrUrArGrArUrGrCrUrCrArArCrCrArA

LvNodal MASO TGCATGGTTAAAAGTCCTTAGAGAT

LvLefty MASO TGCATGGTTAAAAGTCCTTAGAGAT

### Membrane GFP fluorescence experiment

A control construct was built containing a transmembrane sequence in frame with-GFP (Saunders and McClay, 2014). *Membrane-gfp* mRNA was injected into freshly fertilized zygotes at 450ng/µl. A DsiRNA directed against the *membrane-gfp* transcript was injected into one blastomere at the two-cell stage at a concentration of 75.0 µM along with RITC-dextran to mark the injected cell and its progeny. Injected embryos were imaged at multiple timepoints during development. The experiment was conducted more than six times, stopped at many different timepoints, and each experiment imaged up to 10 embryos of more than 100/experiment, all of which had similar fluorescent levels (or lack thereof) at the timepoints tested. The experiment in **Fig. 1** shows one of ≥6 embryos imaged at every timepoint shown.

Controls for non-specific effects of the presence of DsiRNA included a scrambled control designed to the *nodal* sequence used for morpholino knockdowns. Also, a traditional RNAi 18-mer designed to the successful *nodal* DsiRNA sequence was injected to ask if that would work as well (it did not).

ImageJ (Schneider 2012) was used for quantification of GFP fluorescence. A region of interest was marked by hand for each half of the embryo using RITC dextran signal to distinguish between the progeny of each of the 2-cell embryo’s blastomeres. Integrated GFP fluorescence signal per unit area within those regions of interest were measured. A one-sided paired t-test was used to compare change in fluorescent intensity between the two halves of the developing embryos at 4, 6, 8 and 20 hpf. Bonferroni correction was used to adjust for multiple comparisons. Images of 6 embryos were analyzed for each assayed time.

### Experimental approach with DsiRNAs against endogenous genes

Freshly fertilized zygotes are injected with a mixture of DsiRNA + FITC-dextran. DsiRNA reagents are designed and ordered from IDT with design considerations carefully considered as above. Since the requirement for an exact (or almost exact) match with the targeted RNA sequence is necessary, this step is crucial to the success of the experiment. Here, three DsiRNAs were ordered for each gene tested based on the selection sequence described above.

DsiRNAs were reconstituted in RNase free water to a concentration of 100 µM, aliquoted into single use volumes and stored at -80°C until use. (the dilution series reconstituted the DsiRNAs to 400 µM). Injection aliquots are thawed on ice. Injections are performed with final DsiRNA concentrations of 25,50 and 75 µM to optimize concentration for each design, with co-injection of FITC-dextran in 30% glycerol as a tracer. Zygotes are injected using established methods (Cheers and Ettensohn, 2004).

Fluorescent embryos (containing FITC-dextran) are selected and cultured to timepoints where a known, or suspected phenotype should be present if the DsiRNA reagent functions as expected. Some injected embryos are fixed for *in situ* hybridization to determine if there is a visible reduction in RNA relative to control embryos at that stage in the cells of interest. We find that conventional Digoxygenin labeled RNA *in situ* analysis is an excellent indicator of a successful and targeted knockdown, as it matches qPCR and the phenotypes of each of the known perturbations tested. In the experiments, Digoxygenin *in situ* analysis for each gene tested, revealed visual evidence of targeted mRNA knockdown in the expected location in the embryo. Each of the experiments on endogenous genes were performed once to discover functional candidates, then at least 3 times (except for *chat* which repeated in two separate DsiRNA knockdowns performed using eggs from different females). Numbers given in each experiment report quantification of that experiment, and we observed similar outcomes for each repeated experiment.

### RNA In situ analysis

The Protocol for conventional in situ analysis used digoxigenin-11-UTP (Dig) (Roche)-labeled probes (Croce et al., 2003). Hybridization chain reaction (HCR) in situs were performed using probes obtained from Molecular Instruments, Inc. and used according to their protocol.

### qPCR

This protocol was adapted from (Mainland et al., 2017). Total RNA from ∼200 DsiRNA injected, and ∼200 control embryos was extracted and cleaned up using the Direct-zol RNA MicroPrep kit from Zymo Research (Cat no-R2060). mRNA was isolated from total RNA using NEBNext Poly(A) mRNA magnetic isolation module (NEB cat# E7490. Primers (below) were designed to recognize the 5’ half of the mRNA, upstream of the RISC mediated cut. qPCR was performed using QuantiNova Sybr Green RT-PCR kit from Qiagen (Cat # 208152) with Setmar (Histone-lysine N-methyltransferase) and ubiquitin as standards (Hogan et al., 2020).

Primers:

Setmar F – GCCATCATGTCCTTGTCTCA

Setmar R – CACATGAAGCTTGATCAGGTAGTC

Ubiq2 F-CCATAGAGAATGTCAAGGCTAAGA

Ubiq2 R-GGGTTGATTCCTTCTGGATGT

Pks a F-TCGATGATGTTGTGGCTCTATC

Pks a R-GGTGCAGCTTTGGGATTTATG

Nodal a F-CGAGTGGATCATCTACCCAAAG

Nodal a R-GGGTTGTTTGAGTCGGATAAGA

Lefty set F-CATCTCCCAGATCTTACCACTTAC

Lefty set R-GAGAAGAAGCGCAGAGACAA

## Acknowledgements.

We thank other members of our labs for their support and input, and we thank Brennan McDonald and Zach Pracher, enthusiastic undergraduates, for their many questions and encouragement as this project progressed.

---

**Supplementary Fig. S1 Quantification of fluorescence in *mem-gfp,* injected embryos with DsiRNA to *gfp* injected into one cell at the two-cell stage.** Six embryos at each time point imaged were scored in Relative Fluorescent Units (RFU). ** = significance at p<=0.001, and * denoting significance at p<=0.05. The slight increase in fluorescence at 20 hpf is due to migration of fluorescent mesoderm cells into the DsiRNA-injected side in the blastocoel.

**Supplementary Fig. S2**. ***Pks* DsiRNA and localization of *pks1*, *acaca*, and *irf4* (A)** Control *pks1* in situ. (**B**) DsiRNA injected embryo with a low injection volume (based on fluorescent injection tracer) shows a few cells in the embryo (white arrows) that express a low level of *pks1* as seen by *in situ* analysis. (C) A 24 hpf control embryo stained with an HCR *in situ* probe. (**D**) A 24 hpf *pks1* DsiRNA-injected embryo probed with an HCR *in situ* probe. (**E**) A scRNA-seq UMAP of *Lv* development from 2-24 hpf (Massri et al., 2021) showing expression of *pks1* exclusively in a lineage identified as pigment cells. (**F**). Expression of *acaca* (acetyl CoAcarboxylase alpha) on the UMAP plot showing presence in the same cells as *pks1.* (**G**) Expression of *irf4* in a branch of the scRNA-seq UMAP that signature markers identify as ventral mesoderm (blastocoelar cells). Bars = 25µm.

**Supplementary Fig. S3. qPCR results of the DsiRNA knockdowns. (A)** DsiRNA knockdown of *nodal, lefty* and *pks1* showing that each of the three mRNAs is significantly reduced relative to controls at late gastrula. (**B**) DsiRNA knockdown of *lefty, nodal, and pks1* as scored by qPCR of *pks1.* The *lefty* knockdown (too much Nodal) leads to a great reduction of *pks1.* A *nodal* knockdown leads to excess *pks1*, and a repeat of *pks1* knockdown leads to a reduction in *pks1* relative to controls.

**Supplementary Fig. S4. HCR results for *nodal* and *lefty* DsiRNA knockdowns. (A)** control and (**B**) DsiRNA of *nodal.* In the control at the late gastrula stage shown, *nodal* is expressed in the right foregut and right ectoderm. In the DsiRNA knockdown the embryo is radialized and *nodal* fragments are expressed throughout. This reflects an inability of Nodal to establish a ventral community effect, so the mRNA is expressed and fragmented throughout the embryo. (**C**) Control expression of *lefty* in the right foregut and right ectoderm. (**D**) *lefty* fragment expression expands both spatially and quantitatively, since knockdown of *lefty* by DsiRNA leads to an overproduction of Nodal and a corresponding increase in *lefty* transcription. (**E**) Expression of *msp130* is exclusive to skeletogenic cells as seen in a UMAP (Massri et al., 2021). The branch stained by *msp130* reflects the skeletogenic cell lineage based on expression of signature genes. (**F**) Expression of *astacin4* (a zinc metalloproteinase) is seen only in the ventral blastocoelar cell lineage.

**Supplementary Fig. S5. DsiRNA to *chat* mRNA knocks down a late expressed mRNA. (A)** *Chat* is expressed in a subset of neurons of the ciliary band at 24 hpf. **(B)** DsiRNA destroys the *chat* mRNA as seen by digoxygenin *in situ.* **(C., D)** By HCR *chat* expression is seen both in control embryos at 24 hpf (**C),** and in the mRNA fragments of the DsiRNA knockdown **(D).**

## References

Agrawal, N., Dasaradhi, P.V., Mohmmed, A., Malhotra, P., Bhatnagar, R.K., Mukherjee, S.K., 2003. RNA interference: biology, mechanism, and applications. Microbiol Mol Biol Rev 67, 657–685.

Allen, R.L., George, A.N., Miranda, E., Phillips, T.M., Crawford, J.M., Kiehart, D.P., McClay, D.R., 2022. Wound repair in sea urchin larvae involves pigment cells and blastocoelar cells. Dev Biol 491, 56–65.

Amarzguioui, M., Lundberg, P., Cantin, E., Hagstrom, J., Behlke, M.A., Rossi, J.J., 2006. Rational design and in vitro and in vivo delivery of Dicer substrate siRNA. Nat Protoc 1, 508–517.

Ambardekar, V.V., Wakaskar, R.R., Ye, Z., Curran, S.M., McGuire, T.R., Coulter, D.W., Singh, R.K., Vetro, J.A., 2018. Complexation of Chol-DsiRNA in place of Chol-siRNA greatly increases the duration of mRNA suppression by polyplexes of PLL(30)-PEG(5K) in primary murine syngeneic breast tumors after i.v. administration. Int J Pharm 543, 130–138.

Arnone, M.I., Bogarad, L.D., Collazo, A., Kirchhamer, C.V., Cameron, R.A., Rast, J.P., Gregorians, A., Davidson, E.H., 1997. Green Fluorescent Protein in the sea urchin: new experimental approaches to transcriptional regulatory analysis in embryos and larvae. Development 124, 4649–4659.

Bardhan, A., Deiters, A., Ettensohn, C.A., 2021. Conditional gene knockdowns in sea urchins using caged morpholinos. Dev Biol 475, 21–29.

Bolouri, H., Davidson, E.H., 2009. The gene regulatory network basis of the “community effect,” and analysis of a sea urchin embryo example. Dev Biol 340, 170–178.

Britten, R.J., Cetta, A., Davidson, E.H., 1978. The single-copy DNA sequence polymorphism of the sea urchin Strongylocentrotus purpuratus. Cell 15, 1175–1186.

Calestani, C., Rast, J.P., Davidson, E.H., 2003. Isolation of pigment cell specific genes in the sea urchin embryo by differential macroarray screening. Development 130, 4587–4596.

Calestani, C., Rogers, D.J., 2010. Cis-regulatory analysis of the sea urchin pigment cell gene polyketide synthase. Dev Biol 340, 249–255.

Carmell, M.A., Hannon, G.J., 2004. RNase III enzymes and the initiation of gene silencing. Nat Struct Mol Biol 11, 214–218.

Carter, M.G., Sharov, A.A., VanBuren, V., Dudekula, D.B., Carmack, C.E., Nelson, C., Ko, M.S., 2005. Transcript copy number estimation using a mouse whole-genome oligonucleotide microarray. Genome Biol 6, R61.

Cheers, M.S., Ettensohn, C.A., 2004. Rapid microinjection of fertilized eggs. Methods Cell Biol 74, 287–310.

Chendrimada, T.P., Gregory, R.I., Kumaraswamy, E., Norman, J., Cooch, N., Nishikura, K., Shiekhattar, R., 2005. TRBP recruits the Dicer complex to Ago2 for microRNA processing and gene silencing. Nature 436, 740–744.

Coffman, J.A., Wessels, A., Deschiffart, C., Rydlizky, K., 2014. Oral-aboral axis specification in the sea urchin embryo, IV: Hypoxia radializes embryos by preventing the initial spatialization of nodal activity. Dev Biol 386, 302–307.

Croce, J., Lhomond, G., Gache, C., 2003. Coquillette, a sea urchin T-box gene of the Tbx2 subfamily, is expressed asymmetrically along the oral-aboral axis of the embryo and is involved in skeletogenesis. Mech Dev 120, 561–572.

Davidson, E.H., Erwin, D.H., 2006. Gene regulatory networks and the evolution of animal body plans. Science 311, 796–800.

Davidson, E.H., Rast, J.P., Oliveri, P., Ransick, A., Calestani, C., Yuh, C.H., Minokawa, T., Amore, G., Hinman, V., Arenas-Mena, C., et al., 2002. A provisional regulatory gene network for specification of endomesoderm in the sea urchin embryo. Dev Biol 246, 162–190.

Duboc, V., Lapraz, F., Besnardeau, L., Lepage, T., 2008. Lefty acts as an essential modulator of Nodal activity during sea urchin oral-aboral axis formation. Dev Biol 320, 49–59.

Duboc, V., Lapraz, F., Saudemont, A., Bessodes, N., Mekpoh, F., Haillot, E., Quirin, M., Lepage, T., 2010. Nodal and BMP2/4 pattern the mesoderm and endoderm during development of the sea urchin embryo. Development 137, 223–235.

Duboc, V., Rottinger, E., Besnardeau, L., Lepage, T., 2004. Nodal and BMP2/4 signaling organizes the oral-aboral axis of the sea urchin embryo. Dev Cell 6, 397–410.

Duboc, V., Rottinger, E., Lapraz, F., Besnardeau, L., Lepage, T., 2005. Left-right asymmetry in the sea urchin embryo is regulated by nodal signaling on the right side. Dev Cell 9, 147–158.

Fire, A., Albertson, D., Harrison, S.W., Moerman, D.G., 1991. Production of antisense RNA leads to effective and specific inhibition of gene expression in C. elegans muscle. Development 113, 503–514.

Fire, A., Xu, S., Montgomery, M.K., Kostas, S.A., Driver, S.E., Mello, C.C., 1998. Potent and specific genetic interference by double-stranded RNA in Caenorhabditis elegans. Nature 391, 806–811.

Floc’hlay, S., Molina, M.D., Hernandez, C., Haillot, E., Thomas-Chollier, M., Lepage, T., Thieffry, D., 2021. Deciphering and modelling the TGF-beta signalling interplays specifying the dorsal-ventral axis of the sea urchin embryo. Development 148.

Gross, J.M., Peterson, R.E., Wu, S.Y., McClay, D.R., 2003. LvTbx2/3: a T-box family transcription factor involved in formation of the oral/aboral axis of the sea urchin embryo. Development 130, 1989–1999.

Gurdon, J.B., 1988. A community effect in animal development. Nature 336, 772–774.

Hammond, S.M., Caudy, A.A., Hannon, G.J., 2001. Post-transcriptional gene silencing by double-stranded RNA. Nat Rev Genet 2, 110–119.

Hannon, G.J., 2002. RNA interference. Nature 418, 244–251.

Heasman, J., 2002. Morpholino oligos: making sense of antisense? Dev Biol 243, 209–214.

Heasman, J., Kofron, M., Wylie, C., 2000. Beta-catenin signaling activity dissected in the early Xenopus embryo: a novel antisense approach. Dev Biol 222, 124–134.

Hibino, T., Loza-Coll, M., Messier, C., Majeske, A.J., Cohen, A.H., Terwilliger, D.P., Buckley, K.M., Brockton, V., Nair, S.V., Berney, K., et al., 2006. The immune gene repertoire encoded in the purple sea urchin genome. Dev Biol 300, 349–365.

Hogan, J.D., Keenan, J.L., Luo, L., Ibn-Salem, J., Lamba, A., Schatzberg, D., Piacentino, M.L., Zuch, D.T., Core, A.B., Blumberg, C., et al., 2020. The developmental transcriptome for Lytechinus variegatus exhibits temporally punctuated gene expression changes. Dev Biol 460, 139–154.

Howard, E.W., Newman, L.A., Oleksyn, D.W., Angerer, R.C., Angerer, L.M., 2001. SpKrl: a direct target of beta-catenin regulation required for endoderm differentiation in sea urchin embryos. Development 128, 365–375.

Khor, J.M., Ettensohn, C.A., 2023. An optimized Tet-On system for conditional control of gene expression in sea urchins. Development 150.

Kim, D.H., Behlke, M.A., Rose, S.D., Chang, M.S., Choi, S., Rossi, J.J., 2005. Synthetic dsRNA Dicer substrates enhance RNAi potency and efficacy. Nat Biotechnol 23, 222–226.

Konrad, K.D., Arnott, M., Testa, M., Suarez, S., Song, J.L., 2023. microRNA-124 directly suppresses Nodal and Notch to regulate mesodermal development. Dev Biol 502, 50–62.

Lapraz, F., Besnardeau, L., Lepage, T., 2009. Patterning of the dorsal-ventral axis in echinoderms: insights into the evolution of the BMP-chordin signaling network.. PLoS Biol. 7, e1000248.

Lee, R.C., Feinbaum, R.L., Ambros, V., 1993. The C. elegans heterochronic gene lin-4 encodes small RNAs with antisense complementarity to lin-14. Cell 75, 843–854.

Madamanchi, A., Mullins, M.C., Umulis, D.M., 2021. Diversity and robustness of bone morphogenetic protein pattern formation. Development 148.

Mainland, R.L., Lyons, T.A., Ruth, M.M., Kramer, J.M., 2017. Optimal RNA isolation method and primer design to detect gene knockdown by qPCR when validating Drosophila transgenic RNAi lines. BMC Res Notes 10, 647.

Martik, M.L., Lyons, D.C., McClay, D.R., 2016. Developmental gene regulatory networks in sea urchins and what we can learn from them. F1000Res 5.

Massri, A.J., Greenstreet, L., Afanassiev, A., Berrio, A., Wray, G.A., Schiebinger, G., McClay, D.R., 2021. Developmental single-cell transcriptomics in the Lytechinus variegatus sea urchin embryo. Development 148.

Massri, A.J., McDonald, B., Wray, G.A., McClay, D.R., 2023. Feedback circuits are numerous in embryonic gene regulatory networks and offer a stabilizing influence on evolution of those networks. Evodevo 14, 10.

Materna, S.C., Ransick, A., Li, E., Davidson, E.H., 2013. Diversification of oral and aboral mesodermal regulatory states in pregastrular sea urchin embryos. Dev Biol 375, 92–104.

McClay, D.R., 2011. Evolutionary crossroads in developmental biology: sea urchins. Development 138, 2639–2648.

McClay, D.R., Croce, J.C., Warner, J.F., 2021. Conditional specification of endomesoderm. Cells Dev 167, 203716.

Milo, R., and Phillips, R., 2015. Cell Biology by the Numbers. GarlandScience.

Molina, M.D., de Croze, N., Haillot, E., Lepage, T., 2013. Nodal: master and commander of the dorsal-ventral and left-right axes in the sea urchin embryo. Curr Opin Genet Dev 23, 445–453.

Molina, M.D., Lepage, T., 2020. Maternal factors regulating symmetry breaking and dorsal-ventral axis formation in the sea urchin embryo. Curr Top Dev Biol 140, 283–316.

Nam, J., Su, Y.H., Lee, P.Y., Robertson, A.J., Coffman, J.A., Davidson, E.H., 2007. Cis-regulatory control of the nodal gene, initiator of the sea urchin oral ectoderm gene network. Dev Biol 306, 860–869.

Napoli, C., Lemieux, C., Jorgensen, R., 1990. Introduction of a Chimeric Chalcone Synthase Gene into Petunia Results in Reversible Co-Suppression of Homologous Genes in trans. Plant Cell 2, 279–289.

Oulad-Abdelghani, M., Chazaud, C., Bouillet, P., Mattei, M.G., Dolle, P., Chambon, P., 1998. Stra3/lefty, a retinoic acid-inducible novel member of the transforming growth factor-beta superfamily. Int J Dev Biol 42, 23–32.

Oulhen, N., Wessel, G.M., 2016. Albinism as a visual, in vivo guide for CRISPR/Cas9 functionality in the sea urchin embryo. Molecular Reproduction and Development 83:1046–1047.

Peter, I.S., Davidson, E.H., 2015. Genomic Control Process. Development and Evolution. Academic Press, San Diego

Piacentino, M.L., Ramachandran, J., Bradham, C.A., 2015. Late Alk4/5/7 signaling is required for anterior skeletal patterning in sea urchin embryos. Development 142, 943–952.

Pieplow, A., Dastaw, M., Sakuma, T., Sakamoto, N., Yamamoto, T., Yajima, M., Oulhen, N., Wessel, G.M., 2021. CRISPR-Cas9 editing of non-coding genomic loci as a means of controlling gene expression in the sea urchin. Dev Biol 472, 85–97.

Range, R., Lapraz, F., Quirin, M., Marro, S., Besnardeau, L., Lepage, T., 2007. Cis-regulatory analysis of nodal and maternal control of dorsal-ventral axis formation by Univin, a TGF-beta related to Vg1. Development 134, 3649–3664.

Sampilo, N.F., Stepicheva, N.A., Zaidi, S.A.M., Wang, L., Wu, W., Wikramanayake, A., Song, J.L., 2018. Inhibition of microRNA suppression of Dishevelled results in Wnt pathway-associated developmental defects in sea urchin. Development 145.

Saunders, L.R., McClay, D.R., 2014. Sub-circuits of a gene regulatory network control a developmental epithelial-mesenchymal transition. Development 141, 1503–1513.

Schneider, C. A., Rasband, W. S., & Eliceiri, K. W. (2012). NIH Image to ImageJ: 25 years of image analysis. Nature Methods, 9(7), 671–675.

Schulte-Merker, S., Stainier, D.Y., 2014. Out with the old, in with the new: reassessing morpholino knockdowns in light of genome editing technology. Development 141, 3103–3104.

Shabalina, S.A., Koonin, E.V., 2008. Origins and evolution of eukaryotic RNA interference. Trends Ecol Evol 23, 578–587.

Sharp, P.A., 2001. RNA interference--2001. Genes Dev 15, 485–490.

Slota, L.A., McClay, D.R., 2018. Identification of neural transcription factors required for the differentiation of three neuronal subtypes in the sea urchin embryo. Dev Biol 435, 138–149.

Sodergren, E., Weinstock, G.M., Davidson, E.H., Cameron, R.A., Gibbs, R.A., Angerer, R.C., Angerer, L.M., Arnone, M.I., Burgess, D.R., Burke, R.D., et al., 2006. The genome of the sea urchin Strongylocentrotus purpuratus. Science 314, 941–952.

Song, J.L., Stoeckius, M., Maaskola, J., Friedlander, M., Stepicheva, N., Juliano, C., Lebedeva, S., Thompson, W., Rajewsky, N., Wessel, G.M., 2012. Select microRNAs are essential for early development in the sea urchin. Dev Biol 362, 104–113.

Song, J.L., Wessel, G.M., 2007. Genes involved in the RNA interference pathway are differentially expressed during sea urchin development. Dev Dyn 236, 3180–3190.

Steuerwald, N., Cohen, J., Herrera, R.J., Brenner, C.A., 2000. Quantification of mRNA in single oocytes and embryos by real-time rapid cycle fluorescence monitored RT-PCR. Mol Hum Reprod 6, 448–453.

Su, Y.H., Li, E., Geiss, G.K., Longabaugh, W.J., Kramer, A., Davidson, E.H., 2009. A perturbation model of the gene regulatory network for oral and aboral ectoderm specification in the sea urchin embryo. Dev Biol 329, 410–421.

Summerton, J., Weller, D., 1997. Morpholino antisense oligomers: design, preparation, and properties. Antisense Nucleic Acid Drug Dev 7, 187–195.

Yaguchi, S., Taniguchi, Y., Suzuki, H., Kamata, M., Yaguchi, J., 2022. Planktonic sea urchin larvae change their swimming direction in response to strong photoirradiation. PLoS Genet 18, e1010033.

Yaguchi, S., Yaguchi, J., Angerer, R.C., Angerer, L.M., Burke, R.D., 2010. TGFbeta signaling positions the ciliary band and patterns neurons in the sea urchin embryo. Dev Biol 347, 71–81.

Zamore, P.D., Tuschl, T., Sharp, P.A., Bartel, D.P., 2000. RNAi: double-stranded RNA directs the ATP-dependent cleavage of mRNA at 21 to 23 nucleotide intervals. Cell 101, 25–33.

