## Supplemental Figures for "An RNA interference approach for functional studies in the sea urchin and its use in analysis of Nodal signaling gradients"

**Supplementary Fig. S5. DsiRNA to *chat* mRNA knocks down a late expressed mRNA. (A)** *Chat* is expressed in a subset of neurons of the ciliary band at 24 hpf. **(B)** DsiRNA destroys the *chat* mRNA as seen by digoxygenin *in situ.* **(C., D)** By HCR *chat* expression is seen both in control embryos at 24 hpf (**C**), and in the mRNA fragments of the DsiRNA knockdown **(D).**

Fig. S1

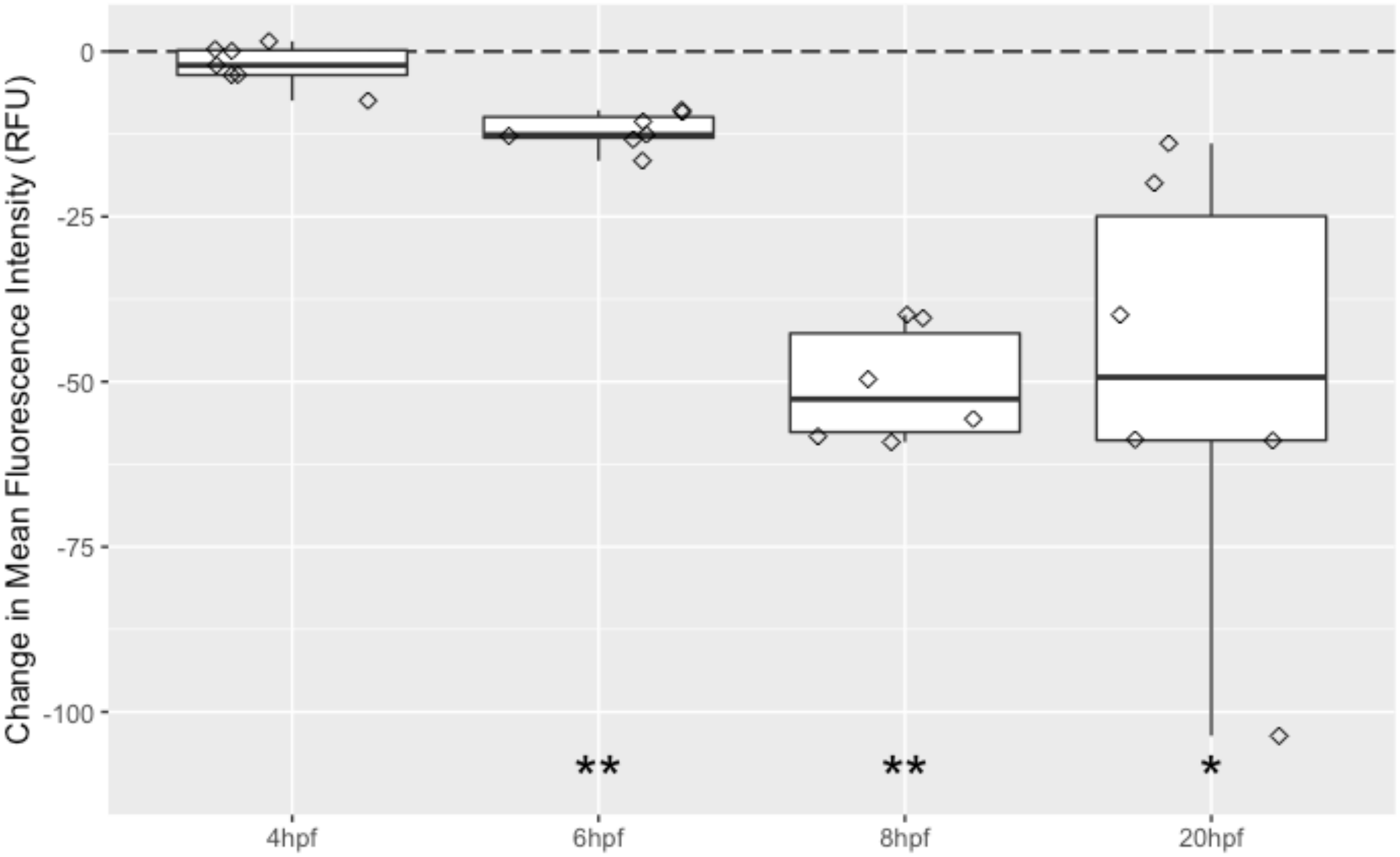

Change in Mean Fluorescence Intensity (RFU)
0
-25
-50
-75
-100
**
**
*
4hpf
6hpf
8hpf
20hpf

Fig. S2

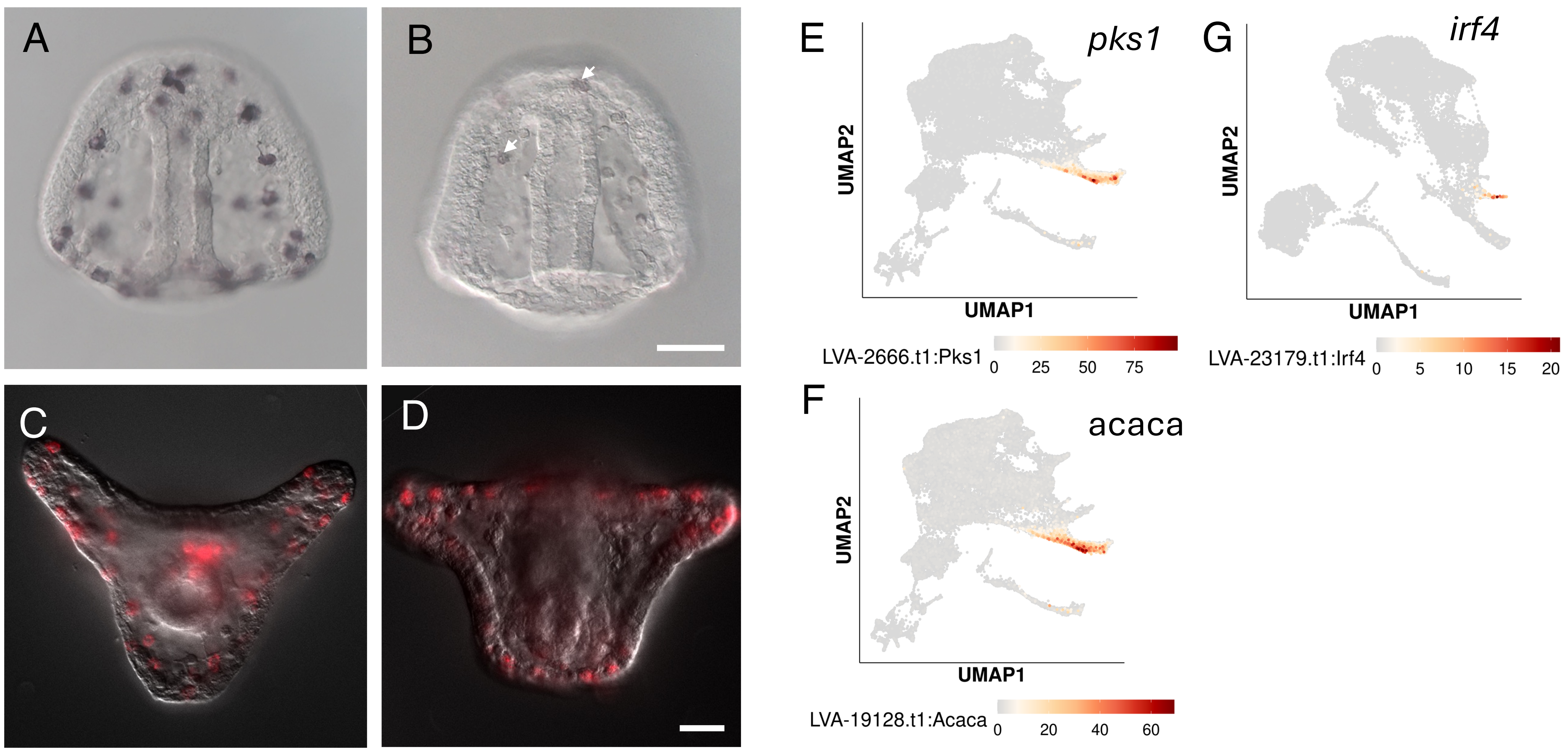
A
B
C
D
E
pks1
UMAP2
UMAP1
LVA-2666.t1:Pks1
0
25
50
75
F
acaca
UMAP2
UMAP1
LVA-19128.t1:Acaca
0
20
40
60
G
irf4
UMAP2
UMAP1
LVA-23179.t1:Irf4
0
5
10
15
20

Fig. S3

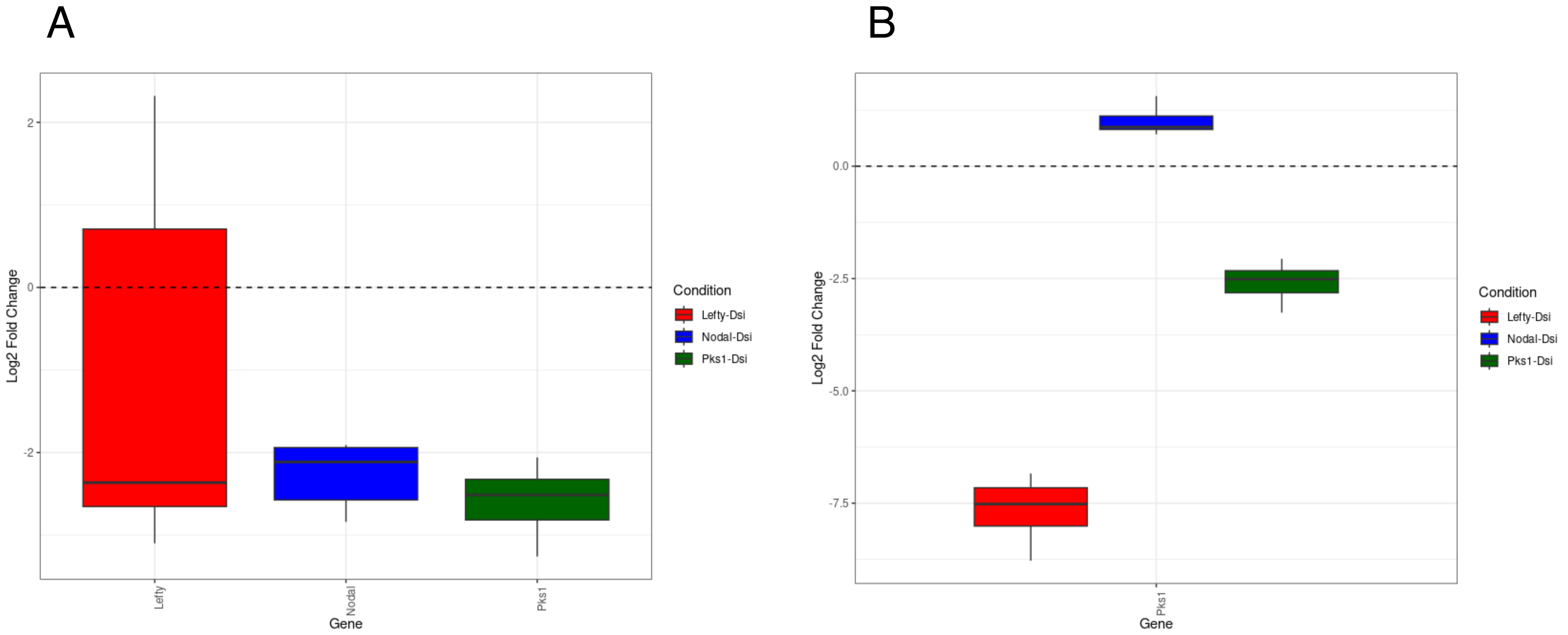
A
B
2
0
-2
Log2 Fold Change
Lefty
Nodal
Pks1
Gene
Condition
Lefty-Dsi
Nodal-Dsi
Pks1-Dsi
0.0
-2.5
-5.0
-7.5
Log2 Fold Change
Pks1
Gene
Condition
Lefty-Dsi
Nodal-Dsi
Pks1-Dsi

Fig. S4

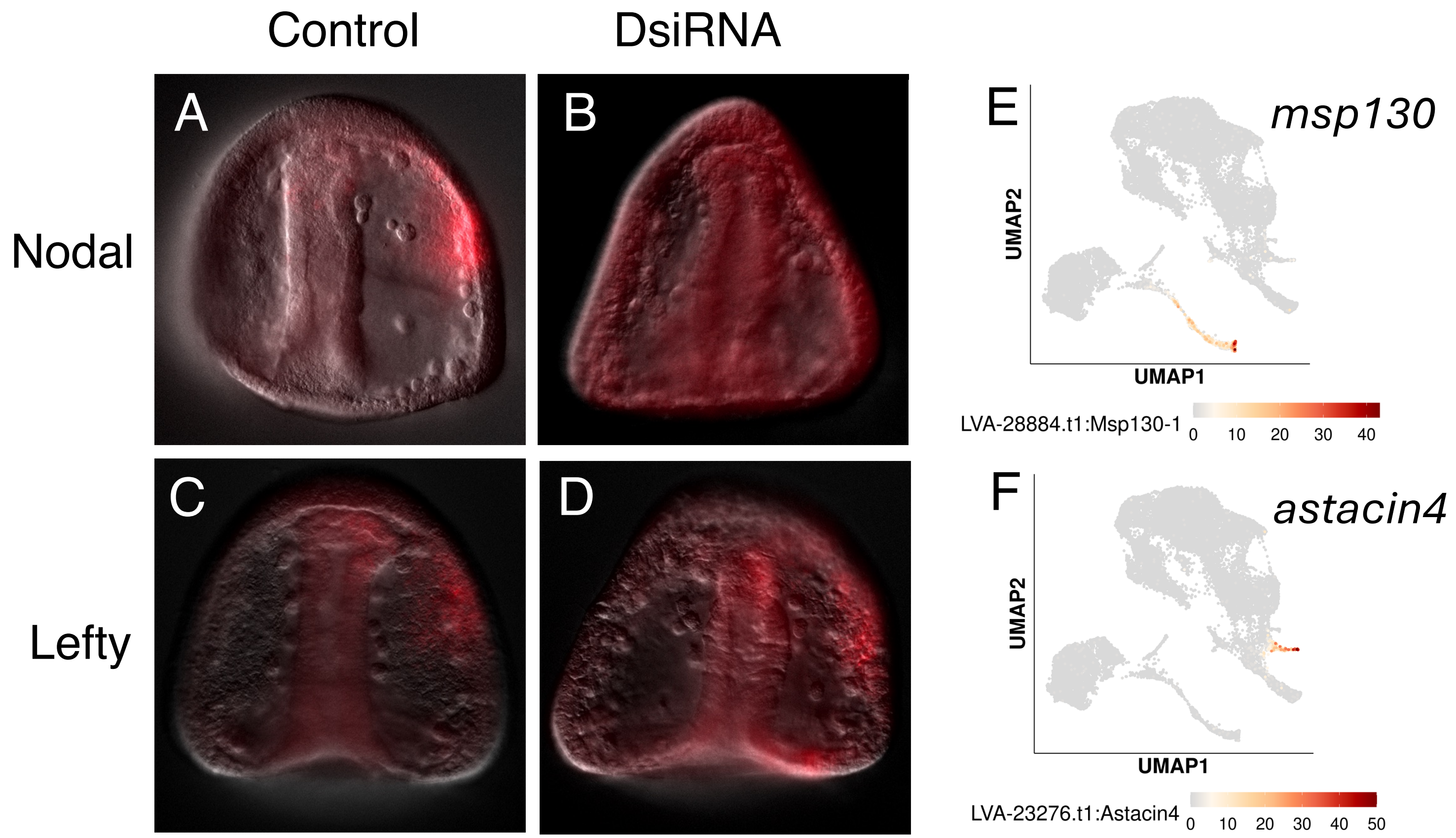

Control
DsiRNA
Nodal
Lefty
A
B
C
D
E
msp130
UMAP2
UMAP1
LVA-28884.t1:Msp130-1
0 10 20 30 40
F
astacin4
UMAP2
UMAP1
LVA-23276.t1:Astacin4
0 10 20 30 40 50

Fig. S5

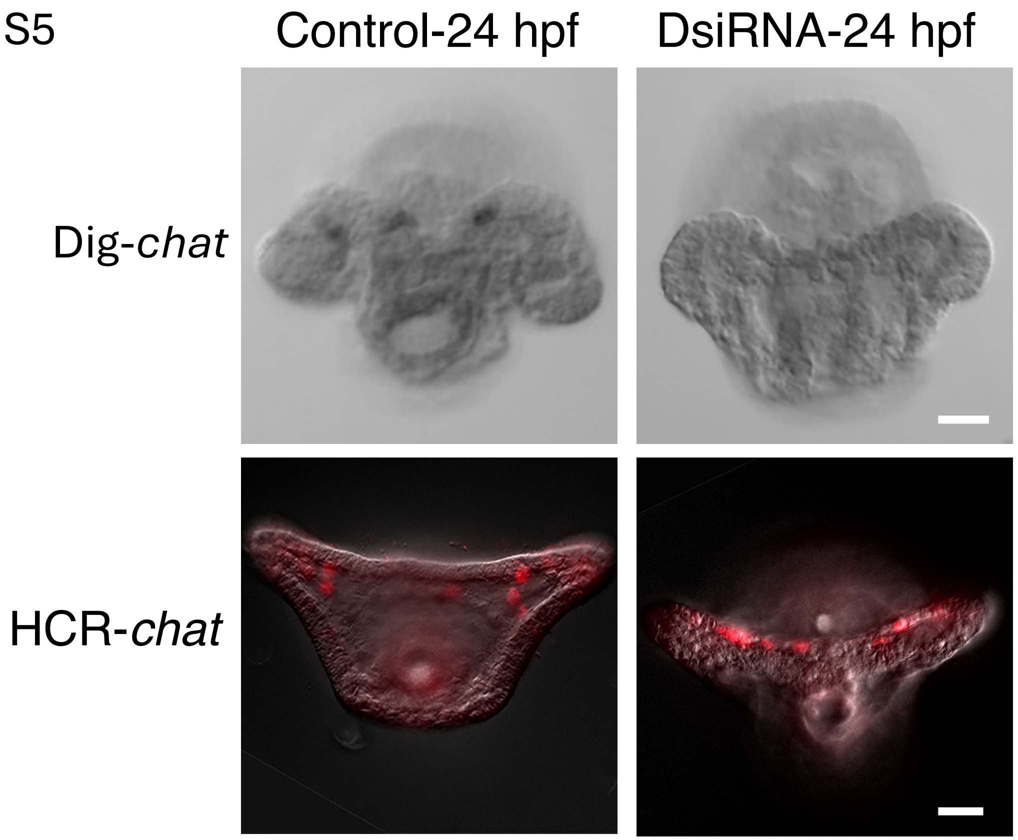
